# Bold but not innovative in an urban exploiter, the red fox (*Vulpes vulpes*)

**DOI:** 10.1101/2022.11.04.515174

**Authors:** F. Blake Morton, Marieke Gartner, Ellie-Mae Norrie, Yacob Haddou, Carl D. Soulsbury, Kristy A. Adaway

**Affiliations:** Department of Psychology, University of Hull, Hull, UK; Atlanta Zoo, Atlanta, Georgia, USA; Institute of Biodiversity, Animal Health and Comparative Medicine, University of Glasgow, Glasgow, UK; School of Life and Environmental Sciences, University of Lincoln, Lincoln, UK

**Keywords:** behavioural flexibility, boldness, human-wildlife conflict, neophobia, problem-solving, urbanisation

## Abstract

Urbanisation is the fastest form of landscape transformation on the planet, but researchers’ understanding of the relationships between urbanisation and animal adaptability is still in its infancy. In terms of foraging, bold and innovative behaviours are proposed to help urban animals access, utilise, and exploit novel anthropogenic food sources. Red foxes (*Vulpes vulpes*) are one of the best known and widespread urban-dwelling species. However, despite frequent stories, images, and videos portraying them as “pests” due to their exploitation of food-related objects (e.g., raiding the contents of outdoor bins), it is unknown whether they are bolder and more innovative in terms of their likelihood of exploiting these resources compared to rural populations. In the current study, we gave novel food-related objects to foxes from 104 locations (one object per location) across a large urban-rural gradient. To access the food, foxes had to use behaviours necessary for exploiting many food-related objects in the real world (e.g., biting, pushing, pulling, or lifting human-made materials). Despite all foxes acknowledging the objects, foxes from 31 locations touched them, while foxes from 12 locations gained access to the food inside. A principal component analysis of urban and other landscape variables (e.g., road, greenspace, and human population density) revealed that urbanisation was significantly and positively related to the likelihood of foxes touching, but not exploiting, the objects. Thus, while urban foxes may be bolder than rural populations in terms of their willingness to physically touch novel food-related objects, our findings are inconsistent with the notion that they are more innovative and pose a general nuisance to people by exploiting these anthropogenic resources.

**Highlights:**

- The impact of urbanisation on animal adaptability remains unclear
- Bold and innovative behaviour may help some urban species thrive
- We studied wild red foxes’ responses to novel food-related objects
- Urban foxes were bolder, but not more innovative, than rural foxes
- Urbanisation may favour bolder, not more innovative, fox behaviour

## Introduction

Urbanisation is the fastest form of landscape transformation on the planet (Angel et al., 2011; Grimm et al., 2008), with 55% of the global human population now living within cities (UN, 2018). Urban environments present wildlife with a range of novel challenges that can include coping with habitat loss and fragmentation (Šálek et al., 2015), increased or novel human disturbances (Rodrigo-Comino et al., 2021), altered competitive interactions (Martin & Bonier, 2018), and new predators or parasites (Guiden et al., 2019; Pedroso-Santos & Costa-Campos, 2020). Species can be characterised based on a gradient of how they adapt to urban environments, including **1)** “*urban avoiders*”, which are restricted to non-urban or remnant natural habitats, **2)** “*urban utilisers*”, which make occasional use of urban areas, and **3)** “*urban dwellers*”, which actively exploit and benefit from urban areas (Fischer et al., 2015). The ability for species to persist and thrive in urban environments is related to a suite of life history, morphological, physiological, behavioural, and cognitive factors (Charmantier et al., 2017; Sol et al., 2014), but researchers’ understanding of how animals adapt to urban environments is still in its infancy.

In terms of foraging, species dwelling in urban areas are likely to encounter novel anthropogenic food sources (Murray et al., 2015; Murray et al., 2018) and behavioural traits, particularly boldness and innovation, are proposed to help urban animals access, utilise, and exploit these resources (Dammhahn et al., 2020; Ducatez et al., 2017; Griffin et al., 2017; Mazza et al., 2021; Mazza & Guenther, 2021). Having a greater tendency to innovate can provide urban wildlife with the behavioural flexibility needed to exploit a wide variety of resources (Reader & Laland, 2003). Being more likely to quickly display such behaviour can enable urban wildlife to exploit these opportunities before they are taken by other animals or removed by city cleaners (Webster et al., 2009). To date, however, not all studies find that urban dwellers are bolder and more innovative for reasons that remain unclear (Griffin et al., 2017; Vincze & Kovacs, 2022).

Red foxes (*Vulpes vulpes*) are one of the best-known and widespread urban-dwelling species (Soulsbury et al., 2010). They are an opportunistic generalist omnivore, which enables them to exploit a diverse range of food items, including mammals, bird, invertebrates, and plants. In urban areas, foxes will also scavenge a wide variety of anthropogenic food items from various sources, including bird feeders, compost heaps, bins, and food provisioned by people (Contesse et al., 2004b; Doncaster et al., 1990; Saunders et al., 1993). Such use of anthropogenic materials suggests that urban foxes are willing to exploit new feeding opportunities, but although urban foxes are often *labelled* as being generally bolder than their rural counterparts, it is unknown whether this is true in all contexts. It is also unknown whether they are more innovative.

Urban foxes often encounter food-related objects that are temporally, physically, and spatially “novel” to them, including **1)** continuous changes to the combination of objects found on streets or in outdoor bins, **2)** objects that look physically different to what animals are accustomed to seeing (e.g., new or modified containers), and **3)** new or familiar objects found in unexpected locations (e.g., randomly discarded trash). Such dynamic changes, combined with frequent encounters, may favour bolder and more innovative behaviour in foxes by enabling them to use new or modified behaviours (i.e., “innovations”) to exploit these resources, particularly shortly after discovering them (e.g., overnight) (Dammhahn et al., 2020; Ducatez et al., 2017; Griffin et al., 2017; Mazza et al., 2021; Mazza & Guenther, 2021). However, despite frequent stories, images, and videos within popular culture portraying urban foxes as “pests” due to their opportunistic foraging behaviour (Schell et al., 2021; Soulsbury & White, 2015), it is unclear whether or to what extent such attitudes are due, in part, to their exploitation of food-related objects, including discarded litter and items found in outdoor bins (Baker et al., 2020; Harris, 1981).

In the past, studies have given novel objects to urban foxes (Padovani et al., 2021), but the objects did not contain food and comparisons with rural populations were not made, making it impossible to evaluate the likelihood of urban foxes behaving bolder and more innovative within this context. Although urban foxes may be more likely to consume novel bait (Gil-Fernandez et al., 2020), this does not necessarily reflect how animals react to other forms of novelty, including human-made objects (Miller et al., 2022). Hence, the current study had two aims: First, to test whether urban foxes are bolder and more innovative than rural populations in terms of exploiting novel food-related objects, and second, to test whether urban foxes are indeed a general nuisance to people because they exploit these anthropogenic resources.

## Methods and materials

### Ethical statement

This study was ethically approved by the Animal Welfare Ethics Board of the University of Hull (FHS356), and was carried out in accordance with guidelines outlined by the Association for the Study of Animal Behaviour (ASAB, 2020). No foxes were handled, all trail cameras were placed away from footpaths to minimise public disturbance, and food items used to attract foxes were not harmful if ingested by other animals, including outdoor pets.

### Study sites and subjects

We studied 200 locations throughout Scotland and England (Figure 1), including areas in and around different cities (e.g., London, Glasgow, Edinburgh, Stirling, Leeds, Hull, Lincoln, Sheffield, and York). These locations covered a wide variety of landscapes, including recreational parks, private gardens, tree plantations, meadows, mixed woodland, coastal and mountainous scrubland, and farmland. Foxes were unmarked and their participation in the study was entirely voluntary. We gained access to 162 of these locations by contacting city councils and other organisations that owned land. The remaining 38 locations were private gardens, which we accessed by advertising the study through Twitter and regional wildlife groups. Our criteria for including any location in the study included: **1)** landowner permission, **2)** accessibility to foxes (e.g., no barriers/fences), **3)** ability to place our equipment out of public view to avoid theft or vandalism, and **4)** the location could not be ≤ 3.5 km from another study area. This latter criterion was used to reduce the chances of sampling the same fox across more than one location because ≥ 3.5 km is larger than the typical dispersal distance and home range diameter of British foxes (Soulsbury et al., 2011; Trewhella et al., 1988). We did not have prior knowledge of fox presence before contacting landowners, and we included locations in the study regardless of whether landowners said foxes lived on or near their land.

**Figure 1:**
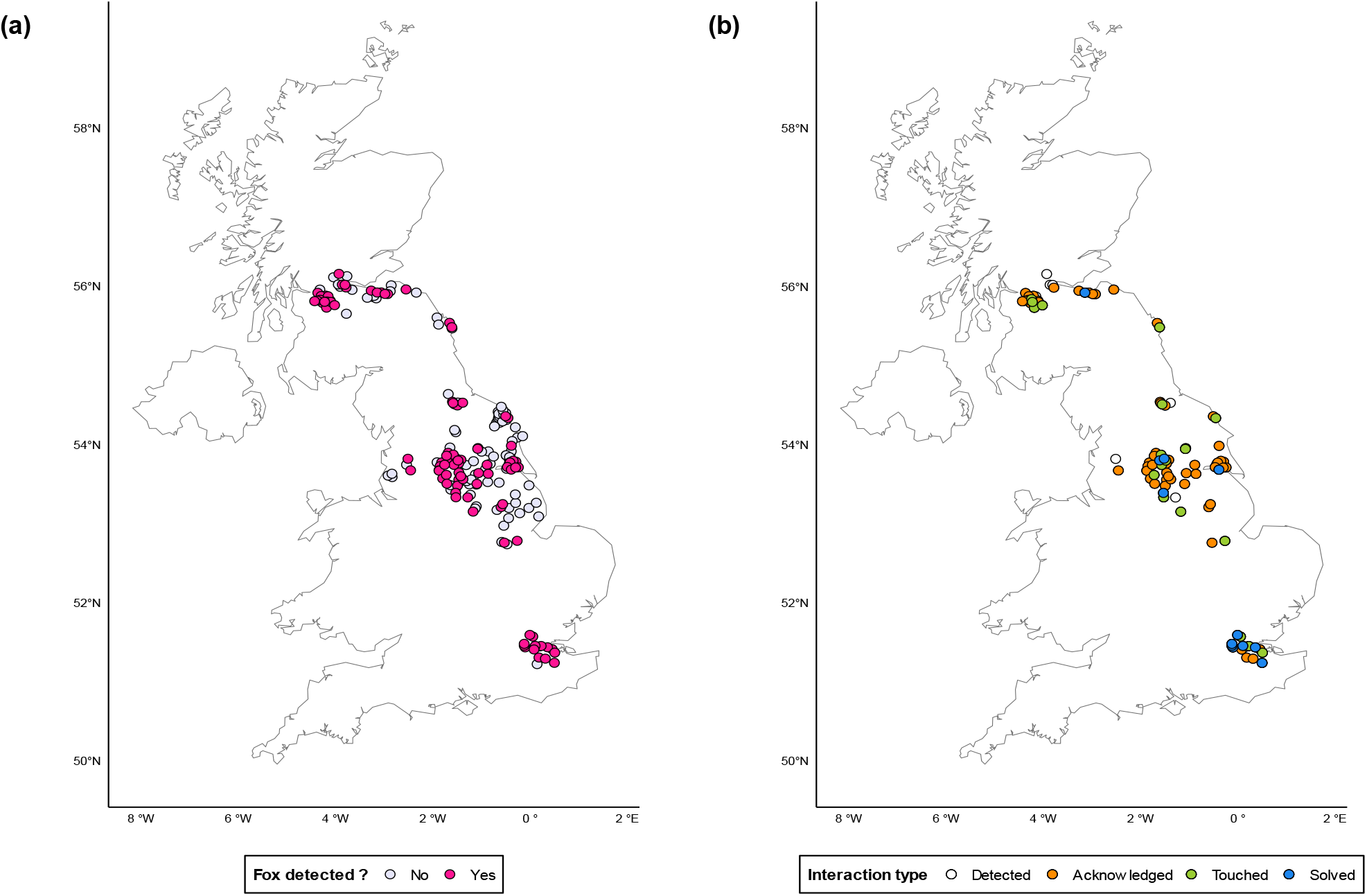
Distribution of locations where objects were deployed across Scotland and England, with (a) foxes detected (yes or no) and (b) whether foxes acknowledged, touched, or exploited (“solved”) the food-related objects.

### Designs and method of administering novel food-related objects

We administered 8 types of food-related objects (Figure 2) across our study locations between August 2021 and November 2023. Only a single object was administered per location, and they were available to foxes for 15.5 ± 1.64 days before we removed them. Although foxes might, of course, respond differently to food-related objects that are left for longer, two weeks is a very typical timeframe for many food-related objects available to British urban foxes (e.g., regular street cleaning and bin services every 1-2 weeks).

**Figure 2:**
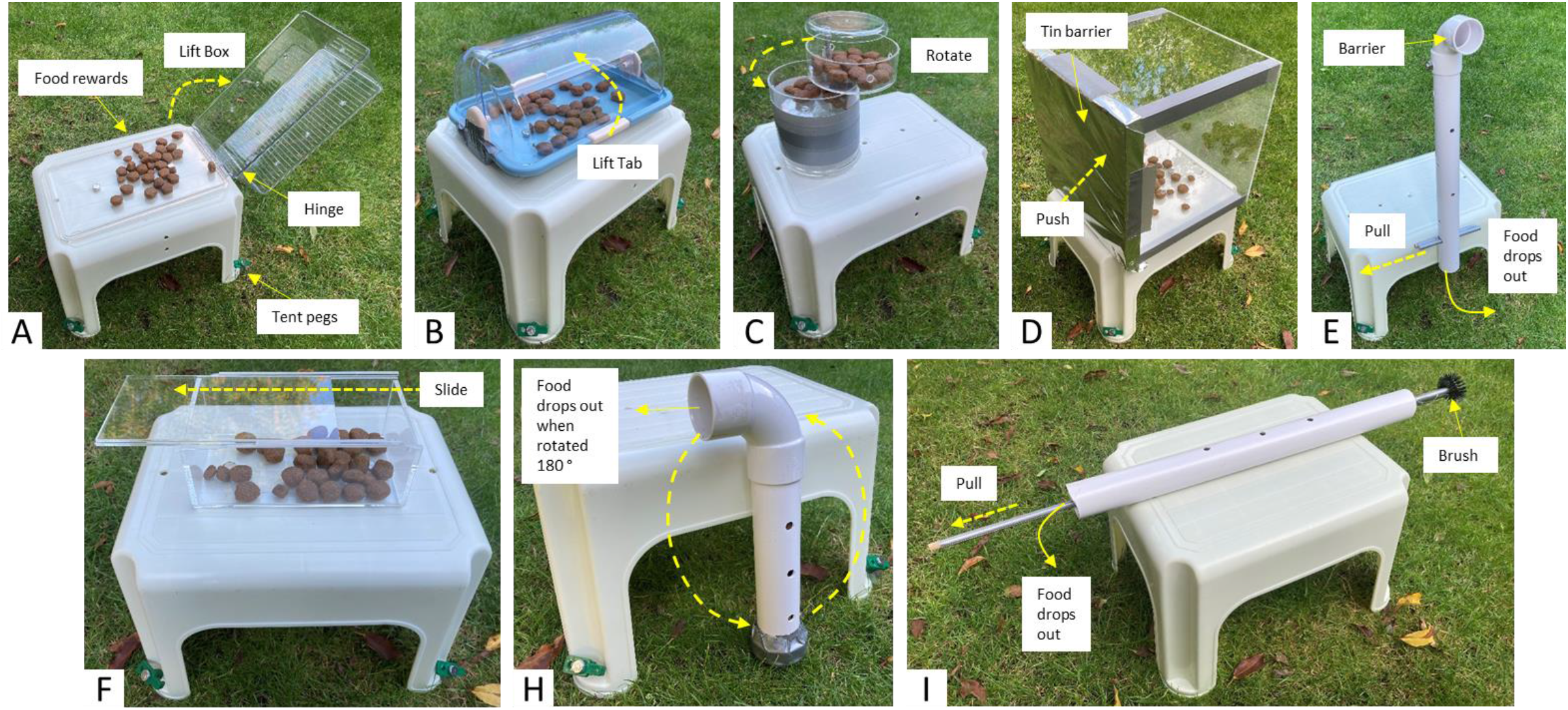
Food-related objects administered to foxes. Yellow dashed arrows indicate the direction of each behaviour needed to retrieve the food rewards inside. Task G was never deployed in the field and hence not depicted in this figure.

The objects were made from basic household materials (e.g., PVC piping, metal screws, and wooden rods). Objects varied in terms of design and materials to ensure that our data on foxes’ behavioural responses were more generalisable and not specific to just one type of object. Objects were “novel” in terms of their location, which we verified by searching for similar objects within the surrounding areas. Objects were also novel in terms of their design, which we assembled ourselves using a unique combination of materials to create objects that are not widely commercially available, making it highly unlikely that foxes would have seen those specific combinations before. ach object had a single ‘free food’ and ‘reward’ condition (Table 1); the ‘free food’ was scattered approximately 1m away from each object. We used different types, combinations, and quantities of food to ensure that our data on foxes’ behavioural responses were more generalisable and not specific to any particular food. All objects were anchored to the plastic platform and had holes drilled into them to facilitate odour cues. Tent pegs were used to anchor the platforms to the ground.

Object C had two levels, each containing food. To access the rewards in Object D, foxes simply had to push through the aluminium side of the box. The lid of Object F was fixed in place and could only be opened by sliding it either to the left or right. Object H had a hidden axle to allow 360° rotation. Object I was administered with the stick already inside the pipe; animals merely had to remove the stick using their mouths, which would indirectly rake the food out.

Researchers were not present when foxes visited, and we did not touch or replenish the food to avoid unnecessary disturbance to the objects. Following APHA guidelines, we cleaned objects with antibacterial soap and 70% alcohol wipes after retrieving them to prevent possible pathogen transmission. We then washed and dried them prior to redeployment. Forty-two objects (21%) were sprayed with scent deodoriser to test whether the scent of objects (e.g., human odour) had a significant effect on fox behaviour.

Since foxes were free-ranging and their participation in the study was entirely voluntary, some foxes might have avoided our testing locations. Nevertheless, the goal of this study was to test foxes’ likelihood of being bold and innovative enough to *exploit* the objects within a two-week period, which required them to physically touch the objects (and hence be detected on camera). We therefore based our analysis on foxes that were at least able and willing to visit the locations.

### Recording fox behaviour from trail cameras

At each location, we horizontally placed a ‘no glow’ (940 nm) infrared motion-sensor camera (Apeman H45) approximately 4m away on a tree trunk. Cameras had a 120 ° sensing angle and a triggering distance of 20 m. Video lengths were set to record for 5 min, with a 5 s trigger delay and a 30 s interval in between each video. Camera lenses were sprayed with defogger and, where possible, minor amounts of understory vegetation were removed between the camera and object to ensure optimal visibility.

### Measuring urban-rural differences in bold and innovative behaviour

A myriad of factors can underpin bold and innovative behaviour, which are not necessarily due to any single variable (Griffin et al., 2014; Lee & Moura, 2015; F. B. Morton et al., 2021; Reader & Laland, 2003). Animals, for example, may not use such behaviour to exploit novel objects if they are too afraid or not hungry. Crucially, however, our goal was to determine *whether* (not why) subjects would display bold and innovative behaviour to exploit food-related objects, and so the only way they could do this was by physically engaging with novel objects themselves.

Foxes could gain access to the food rewards through persistence and by using simple behaviours used to exploit human-made objects in the real world (e.g., using their mouth, nose, and/or paws to bite, push, pull, or lift materials). Some of the designs were inspired from studies of behavioural innovation in other species (Morton, 2021; Rossler et al., 2020; Thornton & Samson, 2012; Visalberghi & Limongelli, 1994). As with these other studies, we defined innovation as any behaviour used to operate and successfully gain access to the food inside each novel object (Figure 2).

To determine whether urban foxes were faster to display bold and innovative behaviours, we compared differences in urban and rural foxes’ likelihood of touching and exploiting (at any point) the objects. We defined ‘touching’ objects in terms of foxes pushing, pulling, licking, and/or biting them, or making physical contact with their nose while smelling them. We defined ‘acknowledging’ objects as a fox turning its head to look/smell in the object’s direction. We tested for inter-observer agreement for all behaviours, and there was excellent agreement (k > .75) between K.A. (who coded all videos), F.B.M. (who developed the definitions and trained K.A.), and several independent coders (Tables S2-5).

While indeed there might be alternative ways of measuring how quickly foxes display bold behaviour, such as walking speed or the latency to approach the objects to within a certain body length, as mentioned before, our research question was related to whether foxes were bold enough to *exploit* them, which required them to physically touch the objects regardless of how long it might have taken them to walk up to them. Similarly, while there may be alternative ways of measuring how quickly foxes innovate, such as the amount of time spent operating the task until a solution was found, this was not possible due to occasional camera malfunctions or some of the videos having poor visibility (e.g., fog or raindrops on the lens); thus, it was more practical, and equally fit for purpose, to analyse how likely urban and rural fox populations were to exploit the food rewards as a function of how many days since the objects were discovered.

### Food tests

Foxes are generalist carnivores and should be highly motivated to consume the food rewards in our study (Saunders & Harris, 2000). To confirm this, we revisited 30 of our locations six months later to leave up to three food conditions, one at a time, on the ground without an object:

- *Condition 1:* 30 chicken-flavoured dried dog food pellets
- *Condition 2:* 15 dried dog food pellets, 15 unsalted peanuts, 1 slice of deli chicken, and 5 sprays of 35mL fish oil mixed with 900mL water
- *Condition 3:* 15 dried dog food pellets, 15 unsalted peanuts, 15 mL honey, 15 mL strawberry jam

All of these locations were within the Yorkshire area. We returned every 3 to 7 days for approximately two weeks to either replenish the same condition or replace it with one of the other three conditions until foxes at each location had an opportunity to discover at least one of the food conditions. Since our goal was to determine whether foxes would consume the food items placed within objects, we recorded the following for all fox visits: **1)** whether the food was still visible when the fox arrived, **2)** whether the fox acknowledged the presence of the food by directing its head and/or nose in the exact spot where we left the food, and **3)** whether the fox consumed the food, including food remnants if some of the food was taken beforehand by another species.

### Factors affecting fox detection and behavioural responses to food-related objects

#### Methodological variables

We examined the impact of object type and food conditions (Table S1), because these may have impacted foxes’ motivation to engage with the objects. We examined the effect of the deodoriser spray because the scent of the puzzles could have deterred foxes (e.g., human scent). Since cameras were not always fully operational (e.g., SD cards full or batteries died), we also examined the impact of the amount of time each camera operated (divided by total days deployment time) after objects were acknowledged by foxes.

Rewards were sometimes exploited by rodents and other organisms that were tiny enough to fit through the holes of objects; thus, whenever possible, we kept records of the presence/absence of rewards at the time of foxes’ initial visits since this might have impacted their ability to detect and engage with the objects. This was done two ways: **1)** by taking a photo of the object whenever researchers visited to switch out the camera’s card, and **2)** looking at the trail camera footage to see whether food was still present. Sometimes we could not determine whether food rewards were still present if, for example, the object was opaque, or we did not return to the location before a fox visited, or the camera footage was not clear enough for us to see inside the transparent objects. At 78 locations, we were able to determine whether food rewards were still present at the time of foxes’ initial visits, but since rewards were missing at only 5 (6.5%) of these locations, we omitted this variable from further analysis given the strong homogeneity of the data.

#### Landscape variables

Most UK residents live within cities and produce many millions of tonnes of waste per year, which leads to significant problems with litter (DEFRA, 2022). Thus, as discussed, foxes exposed to relatively higher levels of urbanisation will have greater access to anthropogenic food-related objects. However, there is no single best way to classify an “urban” versus “rural” population of animals given that the characteristics of urbanisation are so multi-faceted. Hence, to allow us to more accurately evaluate the degree of urbanisation likely experienced by foxes across our study locations, we used a range of variables recommended by Mu et al. (2022), including human population density, road and greenspace density, land coverage (e.g., cropland), and species richness. We also included measures of rainfall, temperature, and elevation because, for example, they factor into cropland suitability.

Landscape data extraction was repeated for a series of circular buffers at 3.5 km from the epicentre of each zone in Figure 1. A Digital Elevation Model raster was sourced from the AWS Open Data Terrain Tiles through the *elevatr* package at a 200m^2^ pixel resolution (Hollister et al., 2021). Average daily mean air temperature over the calendar year (in degrees Celsius) and total precipitation over the calendar year (in millimetres) were obtained from the *HadUK-Grid* climate observation dataset for the year 2021, the most recent available data, at a spatial resolution of 1 km^2^ per pixel (Hollis et al., 2019). Human population size data were collected from the *UK gridded population census 2011* at a 1 km^2^ pixel resolution (Reis et al., 2017). Elevation, temperature, rainfall, and human population size were extracted as the mean raster pixel value within each buffer size. Road density (in m/m^2^) within each buffer was computed by sourcing the highway/road class vector layer from OpenStreetMap (*Planet dump*, 2022). Urban greenspace density (in m^2^/m^2^) was obtained from the *Ordinance Survey Greenspace* vector layer (OPENGREESPACE, 2022). We extracted percent coverage of five land cover classes by employing the UK Centre for Hydrology and Ecology Land Cover 2020 product at a 10 m^2^ resolution (C. S. Morton et al., 2021). The raster is composed of uniquely classified pixels according to categories following the UK Biodiversity Action Plan, which we aggregated into five main land cover categories: *urban* (class 20 and 21), *forest* (class 1 and 2), *grassland* (class 4, 5, 6, 7, 9 and 10), *cropland* (class 3), and *wetland* environments (class 8, 11, 12, 13, 14, 15, 16, 17, 18 and 19). Percent coverage was computed by counting how many pixels within each buffer corresponded to each classified land cover and dividing by the total number of pixels in each buffer. Landscape heterogeneity was also quantified as the effective number of distinct land covers present in each buffer and computed as the exponential of the Shannon-Wiener diversity index (Hill’s numbers equivalent for q=1) (Chao et al., 2014; Hill, 1973).

Landscape variables were calculated using the R programming language version *4*.*2*.*0* within the RStudio IDE version *“Prairie Trillium”* (Team, 2002; Team, 2022). Geospatial vectorial operations were processed utilizing the *sf* R package (Pebesma, 2018) while raster extraction employed the *exactextractr* package (Baston, 2021). Data processing was conducted through the use of the *tidyverse* R packages family (Wickham et al., 2019).

### Statistical analyses

To obtain a global measure of urbanisation from each study location, we entered our landscape variables into a principal component analysis (PCA) with varimax rotation (Team, 2002). A scree test and parallel analysis were used to determine the number of components to extract (Horn, 1965; Morton & Altschul, 2019). Item loadings ≥|.4| were defined as salient for the PCA; items with multiple salient loadings were assigned to the component with the highest loading.

We first tested whether our methodological variables (object type, deodoriser, season, camera operation time, and food conditions) impacted the likelihood of (a) a fox being detected or (b) touching the object, using binary logistic regression. We then carried binary generalised linear mixed effects models (GLMMs) to test the effect of urbanisation on fox behaviour. In our first model, we tested whether detecting a fox was related to habitat (PCA1, PCA2, PCA3), with food type included as a random factor. In our second model, we tested whether the fox touching the object was related to habitat (PCA1, PCA2, PCA3). We also included “*camera*” (i.e., the proportion of time the camera operated after objects were acknowledged by foxes) as an additional covariate, and food as a random effect. Finally, for foxes that touched the objects, we tested whether their ability to access the food inside them was related to habitat (PCA1, PCA2, PCA3). Again, we included the variable “*camera*” as an additional covariate, and food as a random effect. All GLMMs were run using the lme4 package (Bates et al., 2015), with the significance of fixed effects in binomial GLMMS tested using Wald *χ*^2^ tests implemented in the ANOVA function of the car package (Fox & Weisberg, 2019).

Chi-square tests, Cohen’s kappa tests, and the PCA were conducted in IBM SPSS (Version 27). All other analyses were conducted in R version 4.1.1 (Team, 2021). All data are provided in Datasets S1 and S2 in the supplementary materials.

## Results

### Food tests

Of the 30 locations where we conducted food tests, foxes were detected at 23 locations, and 17 foxes discovered at least one of the food conditions before other animals exploited them. All of these foxes approached and consumed the food (Table S6 and Video S2).

### Likelihood of foxes touching and exploiting food-related objects

During the period in which objects were deployed, foxes were recorded at 104 (52%) locations (Figure 1a). Out of the 104 locations where foxes were recorded, it was not possible to tell whether foxes acknowledged objects at 8 (7.7%) locations due to poor visibility or camera malfunctions. In all the remaining 96 locations across all habitats, foxes acknowledged the objects. Foxes went on to touch the objects at 31 locations (32%), and of these, 12 (40%, 1 location could not be determined) exploited the food inside objects (Figure 1b).

### Principal component analysis of landscape characteristics

Across our 200 study locations (Figure 1), a PCA of our ecological and urban measures revealed three components and explained 29.23, 29.05, and 14.47% of the variance, respectively (Table 2, Figure S1, Table S7). Component 1 was labelled “Wilderness” because it was characterised by item loadings related to lower levels of cropland and higher levels of natural and remote spaces (e.g., forests, grasslands, and higher elevations). Component 2 was labelled “Urbanisation” because it was characterised by higher levels of human, road, and greenspace densities, but lower levels of cropland. Component 3 was labelled “Biodiversity” because it was characterised by high levels of landscape heterogeneity and wetlands (i.e., an important habitat for many terrestrial and aquatic species).

**Table 2:** Principal component analysis of ecological and urban variables (N = 200 locations).

| Item | Varimax-rotated components |  |  |
| --- | --- | --- | --- |
|  | PC 1 | PC 2 | PC 3 |
| Temperature | **-.811** | .349 | -.009 |
| Rainfall | **.827** | .179 | .11 |
| Elevation | **.883** | -.008 | .167 |
| Human population size | -.238 | **.875** | -.091 |
| Greenspace density | .066 | **.71** | -.031 |
| Road density | -.143 | **.959** | -.068 |
| LCC: Urban | -.227 | **.939** | -.103 |
| LCC: Forest | **.629** | -.148 | -.094 |
| LCC: Grassland | **.773** | -.131 | .267 |
| LCC: Cropland | **-.507** | **-.702** | -.348 |
| LCC: Wetland | -.346 | -.133 | **.744** |
| Landscape heterogeneity 0 | .203 | 0 | **.716** |
| Landscape heterogeneity 1 | .365 | -.031 | **.742** |

*Note.* Salient loadings are in boldface. LCC=Land cover category. PC = principal component.

### Effect of methodological variables on the likelihood of fox detection and behaviour

Fox detection on camera was not significantly affected by object type, food condition, or deodoriser spray (Table 3). Similarly, the likelihood of foxes touching an object was not related to the object used, deodoriser spray or the proportion of time the camera was operational (Table 3). There was no significant effect of food condition on the likelihood of foxes touching objects (Table 3; Figure 3).

**Table 3:** Fixed effects from two binary logistic regression models tested using the likelihood ratio χ^2^. In each, we tested methodological variables and their likely impact on (a) fox detection and (b) foxes’ physical engagement with objects.

| Model | Parameter | Likelihood Ratio $\chi^2$ | d.f. | P |
| --- | --- | --- | --- | --- |
| (a) Fox detection | Object | 7.03 | 7 | 0.426 |
|  | Food | 10.91 | 11 | 0.451 |
|  | Deodoriser | 0.12 | 1 | 0.731 |
| (b) Fox touches object | Object | 3.45 | 7 | 0.578 |
|  | Food | 18.01 | 11 | 0.081 |
|  | Deodoriser | 0.91 | 1 | 0.341 |
|  | Camera | 0.93 | 1 | 0.335 |

*Note.* "Camera" is the proportion of time the camera operated after objects were acknowledged by foxes.

**Figure 3:**
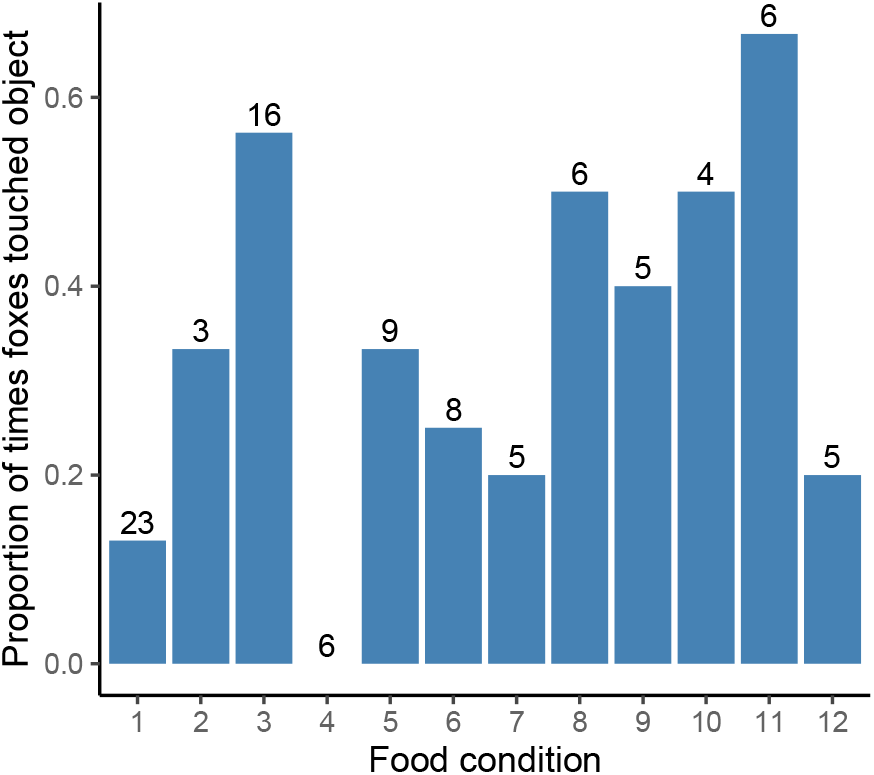
The proportion of times the foxes touched the objects depending on food condition. Numbers over bars indicated the number of times foxes acknowledged the object for each food condition.

### Effect of landscape characteristics on the likelihood of fox detection and touching and exploiting objects

The probability of detecting a fox on camera was significantly lower in more wilderness areas (PCA1: Figure 4a) and greater in more urbanised (PCA2) areas (Table 4; Figure 4b). PCA2 (Urbanisation) was positively associated with foxes touching an object (Table 4; Figure 5), but there was no effect of PCA1 (Wilderness) or PCA3 (Biodiversity) (Table 4). Finally, of those foxes that touched the objects, there was no effect of habitat (PCA1, PCA2, PCA3) on the likelihood of the objects being exploited (Table 4).

**Table 4:** Fixed effects for three binomial GLMM models tested using Wald *χ*^2^ tests. In each, we tested the impact of landscape characteristics on the likelihood of (a) fox detection and foxes (b) touching and (c) exploiting objects.

| Model | Parameter | Wald $\chi^2$ | d.f. | P |
| --- | --- | --- | --- | --- |
| (a) Fox detection | **PCA 1 (Wilderness)** | **6.07** | **1** | **0.014** |
|  | **PCA 2 (Urbanisation)** | **46.33** | **1** | **<0.001** |
|  | PCA 3 (Biodiversity) | 0.29 | 1 | 0.589 |
| (b) Fox touches object | PCA 1 (Wilderness) | 1.47 | 1 | 0.225 |
|  | **PCA 2 (Urbanisation)** | **9.99** | **1** | **0.002** |
|  | PCA 3 (Biodiversity) | 1.48 | 1 | 0.224 |
|  | *Camera* | 0.04 | 1 | 0.844 |
| (c) Fox exploits object | PCA 1 (Wilderness) | 2.04 | 1 | 0.153 |
|  | PCA 2 (Urbanisation) | 1.71 | 1 | 0.191 |
|  | PCA 3 (Biodiversity) | 0.63 | 1 | 0.426 |
|  | *Camera* | 0.15 | 1 | 0.697 |

*Note.* Significant values are in bold. “*Camera*” is the proportion of time the camera operated after objects were acknowledged by foxes.

**Figure 4:**
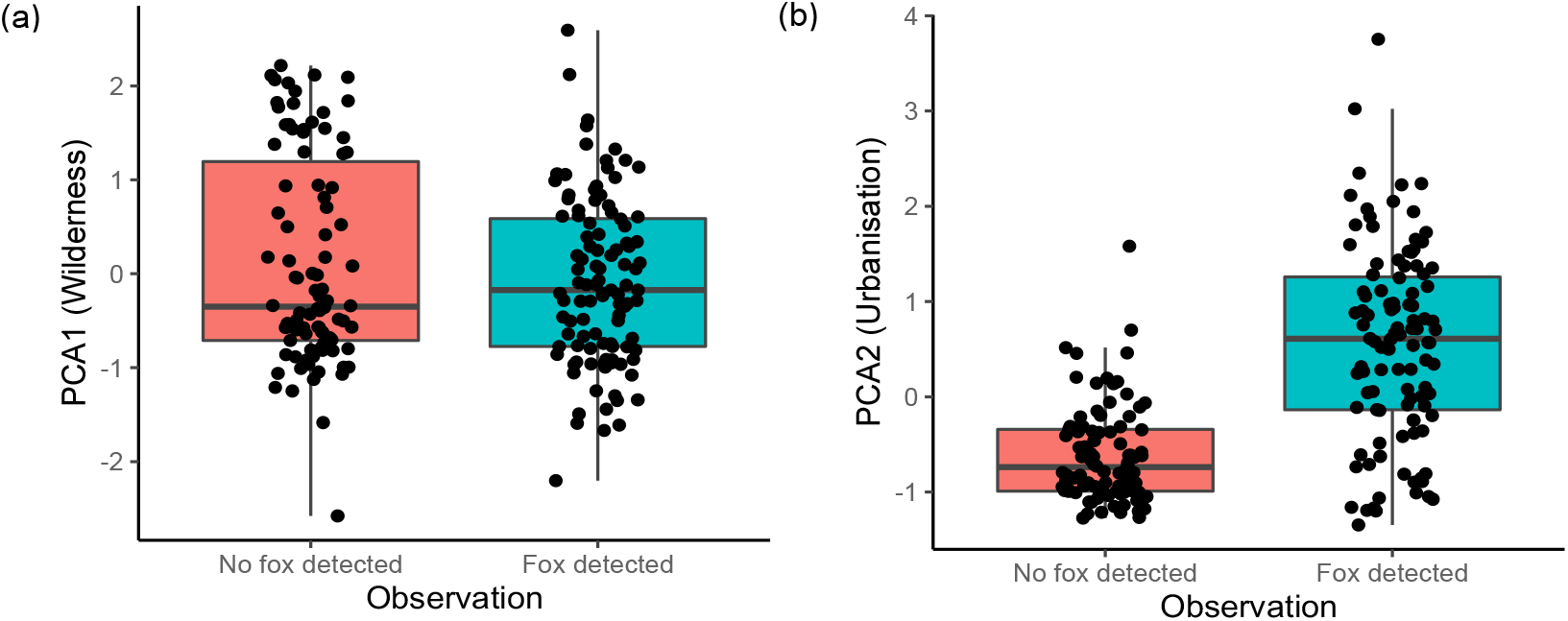
The relationship between (a) foxes being detected by camera in relation to the degree of wilderness (PCA1) and (b) foxes being detected by camera in relation to the degree of urbanisation (PCA2).

**Figure 5:**
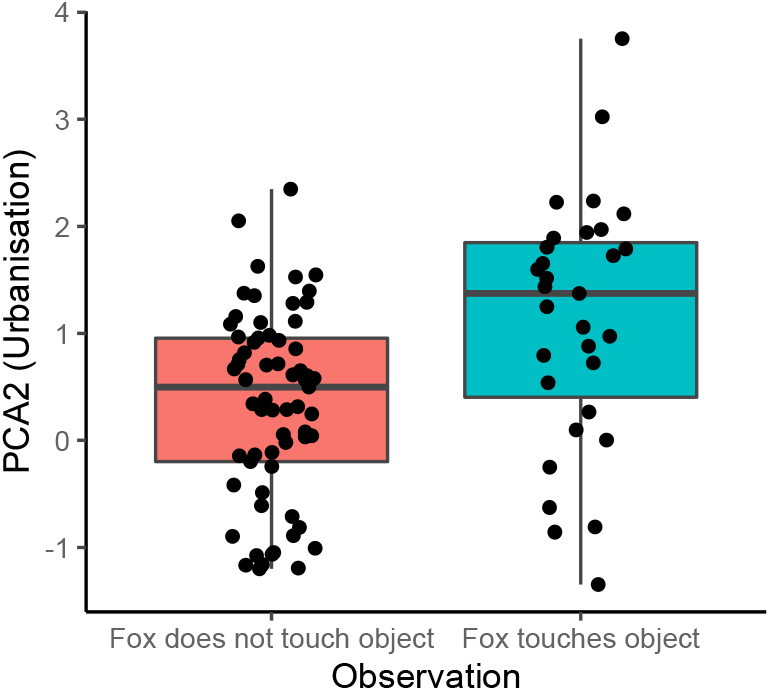
The likelihood of a fox touching a food-related object in relation to the degree of urbanisation (PCA2).

## Discussion

We investigated whether urban foxes are bolder and more innovative than rural populations in terms of exploiting novel food-related objects, and whether such behaviour is consistent with the popular notion that urban foxes are a “pest” because they exploit these anthropogenic resources. Although foxes acknowledged the objects administered in the current study, urbanisation was significantly and positively related to the likelihood of foxes touching, but not exploiting, them. Thus, while urban foxes may be bolder than rural populations in terms of their willingness to physically touch novel food-related objects, our findings are inconsistent with the notion that they are more innovative and pose a general nuisance to people by exploiting them.

Given that we were able to determine the fate of most objects, this rules out the possibility that our cameras significantly missed footage of foxes visiting and exploiting their contents without us knowing it. Foxes always consumed the food rewards when objects were absent despite the presence of a trail camera, ruling out the possibility that the cameras, rather than the food-related objects, were a significant deterrent for them. Since foxes consumed the food rewards when objects were absent, it also rules out the possibility that food-related motivation explains why foxes avoided the objects.

As previously discussed, studies in other species show urban-dwelling animals are more likely than rural populations to physically touch and gain access to novel food-related opportunities (Dammhahn et al., 2020; Ducatez et al., 2017; Griffin et al., 2017; Mazza et al., 2021; Mazza & Guenther, 2021). However, our findings – along with others – illustrate that the relationship between bold and innovative behaviour, particularly with regards to urbanisation, is complex and difficult to generalise across all situations and species (Griffin et al., 2017; Vincze & Kovacs, 2022). Indeed, many other factors likely contribute to whether or how wildlife can adapt to such environments (e.g., dispersal, morphology, and dietary generalist) (Thompson et al., 2021). These studies show, for example, that animals are more innovative in urban environments (field mice, *Apodemus agrarius*; Mazza & Guenther 2021), more innovative in rural environments (spotted hyena: *Crocuta crocuta*; Johnson-Ulrich et al. 2021), or equally innovative in both (this study; house sparrows: *Passer domesticus*; Papp et al. 2014). Thus, for now, although our study suggests that urbanisation may somehow favour (for whatever reason) bolder behaviour in foxes, such behaviour does not necessarily favour them using innovation to exploit food-related opportunities in all contexts (Griffin et al., 2017). Indeed, if that was the case, then more than just 12 (out of 96) urban foxes should have exploited the objects that were administered in the current study after they were discovered.

There are multiple key factors that may separate bold and innovative behaviour. Evidence from birds, at least, suggests that species that are habitat generalists are better at incorporating novel food into their diet, while dietary generalists are more innovative in terms of how they physically acquire food (Ducatez et al., 2015). Red foxes are both habitat and dietary generalists, so it is unclear whether we would predict greater boldness or greater innovation. Our data suggest that boldness is the key behavioural trait; foxes, regardless of location, always consumed food rewards when objects were absent, but not when objects were present. Object neophobia might explain why some foxes avoided the food-related objects (Greggor et al., 2015; Miller et al., 2022; Travaini et al., 2013). Alternatively, given that food resources in urban environments are also very abundant (Ansell, 2005; Contesse et al., 2004b; Harris, 1981), this could explain why urban foxes were motivated to touch, but not necessarily persist and exploit, the unfamiliar food-related objects used in our study. Finally, individual characteristics such as age, sex, dominance, learning speed, and personality might have contributed to fox decision-making and are therefore worth investigating in the future (Fawcett et al., 2017; Griffin et al., 2013; F. B. Morton et al., 2021; Padovani et al., 2021; Soulsbury et al., 2011).

Despite being labelled as a pest, foxes remain a beloved part of urban fauna across the world (Baker & Harris, 2007; Baker et al., 2020; Brand & Baldwin, 2020; Konig, 2008; Nardi et al., 2020), and so future management needs to balance the co-occurrence of both positive and negative human-wildlife interactions within cities (Soulsbury & White, 2015). Our results contrast the UK popular culture’s portrayal of urban foxes as a general ‘nuisance’ because they exploit food-related objects. Such beliefs may stem from specific, highly publicised cases or provocative imagery rather than being typical of urban foxes in general. Indeed, most household surveys (Baker et al., 2004; Harris, 1981), dietary studies (Contesse et al., 2004a), and direct observations (Plumer et al., 2014) show that the image of foxes foraging from bins is uncommon, rather than the norm. Even in our study, most foxes were unlikely to exploit objects when the rewards were relatively large (e.g., 90 dog biscuits). By contrast, our findings from the “free food” condition as well as other studies (Gil-Fernandez et al., 2020) show that when anthropogenic resources are more easily accessible (e.g., no physical barriers), urban foxes may be more likely to exploit such opportunities, which could be due to minimal effort, risk, or both. We suggest that public perceptions of urban foxes stem from their use of freely-available resources, rather than their innovative ability to access unfamiliar resources that require effort.

## Conclusions

Red foxes thrive within urban settings, but contrary to what has been observed in some species, we found that wild urban foxes are, for the most part, no more likely than rural populations to take advantage of novel food-related objects. Thus, while urban foxes may be bolder than rural populations in terms of their willingness to physically touch novel food-related objects, they do not always use innovation to exploit them. The low exploitation rate of food-related objects found in the current study is also contrary to the notion that urban foxes pose a general nuisance to people by exploiting these anthropogenic resources, and therefore calls for a more nuanced view of urban fox behaviour, particularly when it comes to opportunistic foraging.

## Supporting information

Electronic Supplementary Materials

Dataset S1_Fox behaviour

Dataset S2_Principal component analysis

## Acknowledgments

The authors would like to thank everyone involved with the ‘British Carnivore Project’, particularly Sophie Tait, Josh Chatterton, Alice Turner, Eszter Jardan, Dylan Jones, Louise Grunnill, Katherine Sutter, and the citizen scientists who kindly gave us permission to collect data in their gardens. We are very grateful to the many organisations that helped us acquire permission to access other land for the study, particularly staff from the Yorkshire Wildlife Trust, Lincolnshire Wildlife Trust, Scottish Wildlife Trust, The Land Trust, the National Trust, Forestry England (particularly Cath Bashforth), Forestry & Land Scotland, Yorkshire Water, and various city and regional councils (particularly Hull, East Riding of Yorkshire, and Glasgow). Special thanks go to Prof. Hannah Buchanan-Smith (Uni Stirling) and Bruce Gittings (Uni Edinburgh) for their help with various logistics in Scotland, to Dr Alex Weiss (Uni Edinburgh) and Dr Henning Hole (Uni Hull) for statistical discussions, and to Prof. Phyllis Lee (Uni Stirling) and Dr Kevin Parson (Uni Glasgow) for helpful comments on an earlier draft of the manuscript. Finally, FBM is grateful to the University of Hull, UKRI Natural Environment Research Council (NERC) (Grant No. NE/X018342/1), and EU Social Fund Plus for funding.

## Notes

**Conflict of interest statement:** The authors declare no conflict of interest.

### Competing Interest Statement

The authors have declared no competing interest.

### Summary of Updates

New results and additional data collected for the study. Revisions to introduction, methods, and discussion also made.

