## Supplementary material for "Bold but not innovative in an urban exploiter, the red fox (*Vulpes vulpes*)": Electronic Supplementary Materials

Table S1. Food conditions placed 1 metre around ("free food") and inside ("rewards") each novel object.

| **Food Condition** | **Free Food (1 metre around objects)** | **Food Rewards (inside objects)** |
|---|---|---|
| 1 | 15 dried dog food pellets | 30 dried dog food pellets |
| 2 | 30 dried dog food pellets | 15 dried dog food pellets |
| 3 | 15 dried dog food pellets | 90 dried dog food pellets |
| 4 | 90 dried dog food pellets | 15 dried dog food pellets |
| 5 | 15 dried dog food pellets | 15 dried dog food pellets, plus 15 unsalted peanuts (no shells), plus 1 slice of thinly pressed deli chicken, plus *five sprays* of 35mL of fish oil mixed with 900mL water. |
| 6 | 15 dried dog food pellets, plus 15 unsalted peanuts (no shells), plus 1 slice of thinly pressed deli chicken, plus *five sprays* of 35mL of fish oil mixed with 900mL water. | 15 dried dog food pellets |
| 7 | 15 dried dog food pellets | 45 dog food pellets, 45 peanuts, 3 slices of chicken, plus *five sprays* of 105mL of fish oil mixed with 900mL water. |
| 8 | 45 dog food pellets, 45 peanuts, 3 slices of chicken, plus *five sprays* of 105mL of fish oil mixed with 900mL water. | 15 dried dog food pellets |
| 9 | 15 dried dog food pellets | 15 dried dog food pellets plus 15 unsalted peanuts (no shells) mixed with 15mL of honey and 15mL of fruit jam. |
| 10 | 15 dried dog food pellets plus 15 unsalted peanuts (no shells) mixed with 15mL of honey and 15mL of fruit jam. | 15 dried dog food pellets |
| 11 | 45 dried dog food pellets, 45 unsalted peanuts, 45mL of honey and 45 mL of fruit jam | 15 dried dog food pellets |
| 12 | 15 dried dog food pellets | 45 dried dog food pellets, 45 unsalted peanuts, 45mL of honey and 45 mL of fruit jam |

Table S2. Data coded by F.B.M. and two independent observers to test the reliability of coding foxes' acknowledgement of objects.

| Fox video | Acknowledged puzzle? | | |
|---|---|---|---|
| | **FBM** | **Coder 2** | **Coder 3** |
| 1 | 1 | 1 | 1 |
| 2 | 0 | 1 | 0 |
| 3 | 0 | 1 | 1 |
| 4 | 1 | 1 | 1 |
| 5 | 1 | 1 | 1 |
| 6 | 1 | 1 | 1 |
| 7 | 1 | 1 | 1 |
| 8 | 1 | 1 | 1 |
| 9 | 1 | 1 | 1 |
| 10 | 0 | 0 | 0 |
| 11 | 1 | 1 | 1 |
| 12 | 1 | 1 | 1 |
| 13 | 0 | 0 | 0 |
| 14 | 1 | 1 | 1 |
| 15 | 1 | 1 | 1 |
| 16 | 1 | 1 | 1 |
| 17 | 1 | 1 | 1 |
| 18 | 1 | 1 | 1 |
| 19 | 1 | 1 | 1 |
| 20 | 1 | 1 | 1 |
| 21 | 1 | 1 | 1 |
| 22 | 1 | 1 | 1 |
| 23 | 0 | 0 | 0 |
| 24 | 0 | 0 | 0 |

*Note*. 1=behaviour observed, 0=behaviour not observed. Inter-observer reliabilities between F.B.M. and the two independent coders were $k$=.75 and $k$=.88, respectively, for 'acknowledge puzzle'.

Table S3. Data coded by K.A. (who coded all videos) and F.B.M. (who trained K.A.) to ensure K.A. was reliable.

| | Acknowledged puzzle? | |
|---|---|---|
| Fox video | **Coder 1** | **Coder 2** |
| 1 | 1 | 1 |
| 2 | 0 | 0 |
| 3 | 0 | 0 |
| 4 | 1 | 1 |
| 5 | 1 | 1 |
| 6 | 1 | 1 |
| 7 | 1 | 1 |
| 8 | 1 | 1 |
| 9 | 1 | 1 |
| 10 | 0 | 0 |
| 11 | 1 | 1 |
| 12 | 1 | 1 |
| 13 | 1 | 0 |
| 14 | 1 | 1 |
| 15 | 1 | 1 |
| 16 | 1 | 1 |
| 17 | 1 | 1 |
| 18 | 1 | 1 |
| 19 | 1 | 1 |
| 20 | 1 | 1 |
| 21 | 1 | 1 |
| 22 | 1 | 1 |
| 23 | 0 | 0 |
| 24 | 0 | 0 |

*Note*. 1=behaviour observed, 0=behaviour not observed. Inter-observer reliabilities between F.B.M. and K.A. were $k$=.88 for ‘acknowledge puzzle’.

Table S4. Data coded by K.A. at two time periods separated by several months to test for intra-observer consistency.

| | Acknowledged puzzle? | |
|---|---|---|
| Fox video | **Time 0** | **Time 1** |
| 1 | 1 | 1 |
| 2 | 0 | 0 |
| 3 | 1 | 1 |
| 4 | 1 | 1 |
| 5 | 1 | 1 |
| 6 | 1 | 1 |
| 7 | 1 | 1 |
| 8 | 1 | 1 |
| 9 | 1 | 1 |
| 10 | 1 | 1 |
| 11 | 1 | 1 |
| 12 | 1 | 1 |
| 13 | 1 | 0 |
| 14 | 1 | 1 |
| 15 | 0 | 0 |
| 16 | 1 | 1 |
| 17 | 1 | 1 |
| 18 | 1 | 1 |
| 19 | 0 | 0 |
| 20 | 1 | 1 |
| 21 | 1 | 1 |
| 22 | 0 | 0 |

Note. 1=behaviour observed, 0=behaviour not observed. Agreement between K.A.'s scores at Time 0 and 1 were *k*=.86 for 'acknowledge puzzle'.

Table S5. Data coded by K.A. (who coded all the videos) and an independent observer to test the reliability of coding foxes touching and exploiting food-related objects.

| | Fox detected? | | Did a fox touch object within 2 weeks? | | Did a fox exploit object within two weeks? | |
|---|---|---|---|---|---|---|
| | K.A. | Coder 2 | K.A. | Coder 2 | K.A. | Coder 2 |
| 1 | 1 | 1 | 0 | 0 | 0 | 0 |
| 2 | 1 | 1 | 0 | 0 | 0 | 0 |
| 3 | 1 | 1 | 1 | 1 | 1 | 1 |
| 4 | 1 | 1 | 1 | 1 | 1 | 1 |
| 5 | 1 | 1 | 0 | 0 | 0 | 0 |
| 6 | 0 | 0 | | | | |
| 7 | 1 | 1 | 1 | 1 | 0 | 0 |
| 8 | 1 | 1 | 1 | 1 | 0 | 0 |
| 9 | 1 | 1 | 1 | 1 | 0 | 0 |
| 10 | 1 | 1 | 1 | 1 | 1 | 1 |
| 11 | 1 | 1 | 1 | 1 | 1 | 1 |
| 12 | 1 | 1 | 1 | 1 | 0 | 0 |
| 13 | 1 | 1 | 1 | 1 | 1 | 1 |
| 14 | 1 | 1 | 1 | 1 | 1 | 1 |
| 15 | 1 | 1 | 0 | 0 | 0 | 0 |
| 16 | 1 | 1 | 1 | 1 | 1 | 1 |

*Note*. 1=behaviour observed, 0=behaviour not observed. Inter-observer reliabilities between observers were $k$=1 for all behaviours.

Table S6. Food conditions consumed by foxes.

| Location | Condition 1 | Condition 2 | Condition 3 |
|---|---|---|---|
| 1 | Yes | Yes | Yes |
| 2 | --- | Yes | Yes |
| 3 | --- | Yes | --- |
| 4 | Yes | Yes | Yes |
| 5 | Yes | Yes | Yes |
| 6 | Yes | Yes | Yes |
| 7 | --- | Yes | Yes |
| 8 | --- | --- | Yes |
| 9 | --- | Yes | Yes |
| 10 | Yes | Yes | Yes |
| 11 | Yes | Yes | Yes |
| 12 | --- | --- | Yes |
| 13 | --- | Yes | Yes |
| 14 | --- | --- | Yes |
| 15 | --- | Yes | --- |
| 16 | --- | Yes | --- |
| 17 | Yes | Yes | Yes |

*Note*. “---" = food condition was not administered.

Table S7. Random data eigenvalues from parallel analysis to determine number of components to extract for principal component analysis.

| Component | Mean Eigenvalue | Percentile Eigenvalue |
|---|---|---|
| 1 | 1.440556 | 1.532997 |
| 2 | 1.325796 | 1.404585 |
| 3 | 1.243438 | 1.305825 |
| 4 | 1.169502 | 1.219462 |
| 5 | 1.101734 | 1.147657 |
| 6 | 1.043224 | 1.088601 |
| 7 | .982809 | 1.034463 |
| 8 | .928814 | .974348 |
| 9 | .871461 | .908924 |
| 10 | .813521 | .853746 |
| 11 | .756786 | .806806 |
| 12 | .698160 | .754805 |
| 13 | .624200 | .677534 |

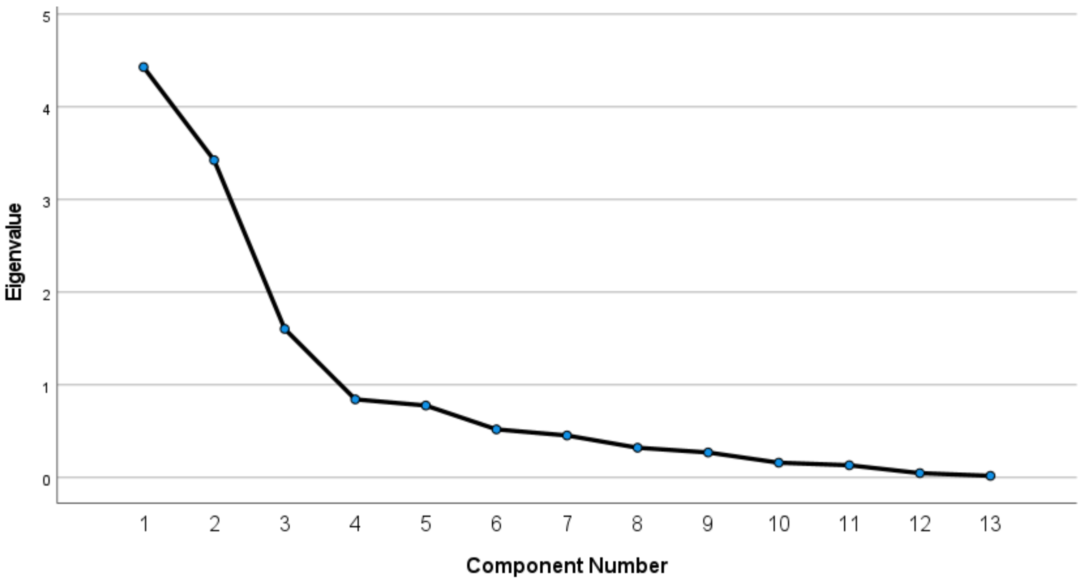


Figure S1. Scree plot for PCA of ecological and urban variables (N = 200 locations).

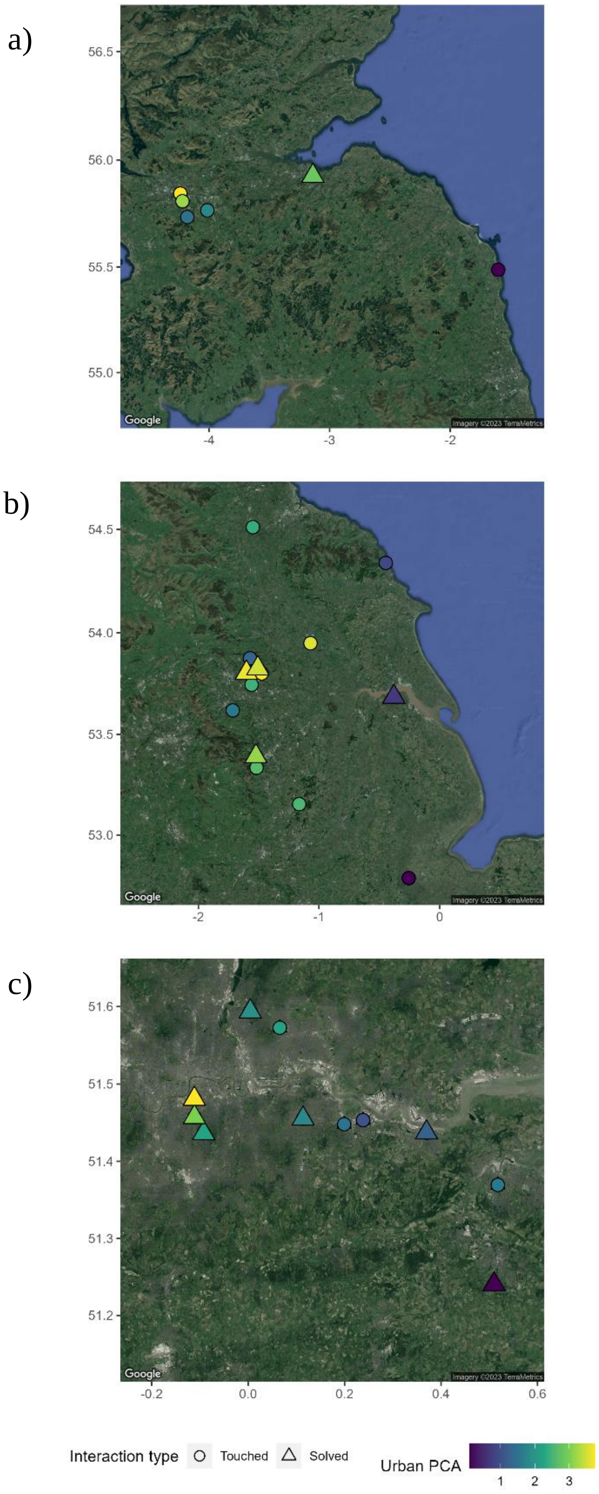


Figure S2. Locations where foxes touched versus exploited (“solved”) food-related objects in relation to the degree of urbanisation (■ = least urban, ■ = most urban), including (a) locations from Scotland and Northumberland, (b) Yorkshire, Lincolnshire, and Nottinghamshire, and (c) London and Kent.

**Supplementary Video Files**

**Link to all videos:** https://youtube.com/playlist?list=PLGW3Yb735jIiADEG8yQtN-6w3pRx1PMqz

**Video S1.** Examples of foxes touching and exploiting objects

**Video S2.** Examples of foxes eating food rewards when objects not present
